# Nodal is a short-range morphogen with activity that spreads through a relay mechanism in human gastruloids

**DOI:** 10.1101/2021.04.14.439902

**Authors:** Lizhong Liu, Anastasiia Nemashkalo, Ji Yoon Jung, Sapna Chhabra, M. Cecilia Guerra, Idse Heemserk, Aryeh Warmflash

## Abstract

Morphogens are signaling molecules that convey positional information and dictate cell fates during development. Little is known about how morphogen gradients are created and interpreted during mammalian embryogenesis. Here we take advantage of a human gastruloid model to visualize endogenous Nodal protein in living cells. We show that Nodal is extremely short range so that Nodal protein is limited to the immediate neighborhood of source cells. Nodal activity spreads through a relay mechanism in which Nodal production induces neighboring cells to transcribe Nodal. We further show that the Nodal inhibitor Lefty, while biochemically capable of long-range diffusion, also acts locally to control the timing of Nodal spread and therefore of mesoderm differentiation during patterning. Our study establishes a novel paradigm for tissue patterning by an activator-inhibitor pair.

## Introduction

One of the central questions in developmental biology is how spatial distributions of extracellular morphogens are controlled and interpreted to produce patterns of cell fates. While it is commonly believed that positional information is conveyed by long-range gradients of morphogens (*1, 2*), direct evidence is difficult to obtain as they are present in the extracellular space and often effective at low (nanomolar) concentrations (*3*). Nodal, a secreted member of the transforming growth factor ß (TGF-ß) family, is essential for stabilization of the epiblast state, induction of both the anterior-posterior axis and left-right axes, and specification of mesoderm and endoderm (*4, 5*). A differential diffusivity (Turing) model has been proposed to account for the creation of a Nodal gradient through interactions with Lefty1 and Lefty2 (hereafter Lefty1/2) that diffuse faster than Nodal (*6*). Evidence from ectopic expression experiments performed in Zebrafish supports this model (*6*–*8*), however, overexpression studies can saturate receptors and other binding sites, altering the spatial range of secreted molecules. Direct validation has been prevented by the difficulty of measuring endogenous morphogens during patterning. In addition to paracrine signaling over several cell diameters, other models which do not rely on extracellular transport, including a relay model in which signaling induces the transcription of the gene encoding the morphogen in neighboring cells have also been suggested (*9*–*13*). While there are six Nodal genes in Xenopus and two in Zebrafish (*6*–*8, 11, 14*), there is only one mammalian Nodal gene, which provides an opportunity to study the ligand distribution for this pathway by studying a single protein. To date, owing to technical limitations, endogenous Nodal protein has never been visualized, therefore its spatiotemporal distribution in mammalian embryos remains a mystery.

Here we visualize and quantitatively measure the levels of endogenous Nodal and Lefty1/2 mRNA and protein in a model of human germ layer patterning, micropatterned human embryonic stem cells (hESCs)-based gastruloids (*15, 16*). In this model, exogenous BMP signaling triggers a cascade of signaling through the BMP, Wnt, and Nodal pathways causing germ layer differentiation. This revealed that Nodal protein and signaling activity are restricted to cells that immediately touch producing cells. Through genetic manipulation, we show that propagation of the Nodal signal requires that the Nodal gene be intact in receiving cells, suggesting a transcriptional relay model. As Nodal activity spreads inwards in the gastruloid through this relay, Lefty is expressed specifically at the signaling front. Deletion of both Lefty genes leads to earlier and more rapid spreading and an expansion of mesendoderm differentiation. This suggests that the function of Lefty during gastrulation is to control the timing and spread of the Nodal signaling wave to properly induce mesendoderm.

## Results and discussion

### Visualization of fully functional endogenous Nodal protein

To visualize endogenous Nodal protein, we generated hESCs with an mCitrine::Nodal (hereafter, cNodal) fusion allele. In these cells, the fluorescent mCitrine is inserted between the Nodal pro-domain and mature domains (Fig. S1). After being translated, the full-length protein will be cleaved by convertases (*17*), splitting the pro-domain from the mCitrine-tagged mature domain. We generated both heterozygous Nodal^+/Cit^ and homozygous Nodal^Cit/Cit^ cell lines. Unless specifically noted otherwise, all results reported are for homozygous Nodal^Cit/Cit^ cells. The Nodal-citrine protein was readily visualized in modified cells where Nodal could be found in the cytoplasm and on the basal side of the cells with less protein localized to the apical side. We also observed occasional long cell protrusions containing Nodal-citrine protein (Fig. S2).

We found that mCitrine tagging does not compromise hESCs pluripotency (Fig. S3), and natural regulation of Nodal expression is preserved (Fig. 1). In particular, cNodal expression level increases in response to addition of Activin, a ligand of Activin/Nodal signaling, and Wnt3A, an upstream signal (Fig. 1A, B). In line with previous studies, BMP4 does not stimulate Nodal expression directly (Fig 1A), however, experiments in gastruloids confirm that BMP signaling induces Nodal through a Wnt intermediate, as with untagged Nodal (Fig 1B, supplementary movies) (*18, 19*).

**Figure 1.**
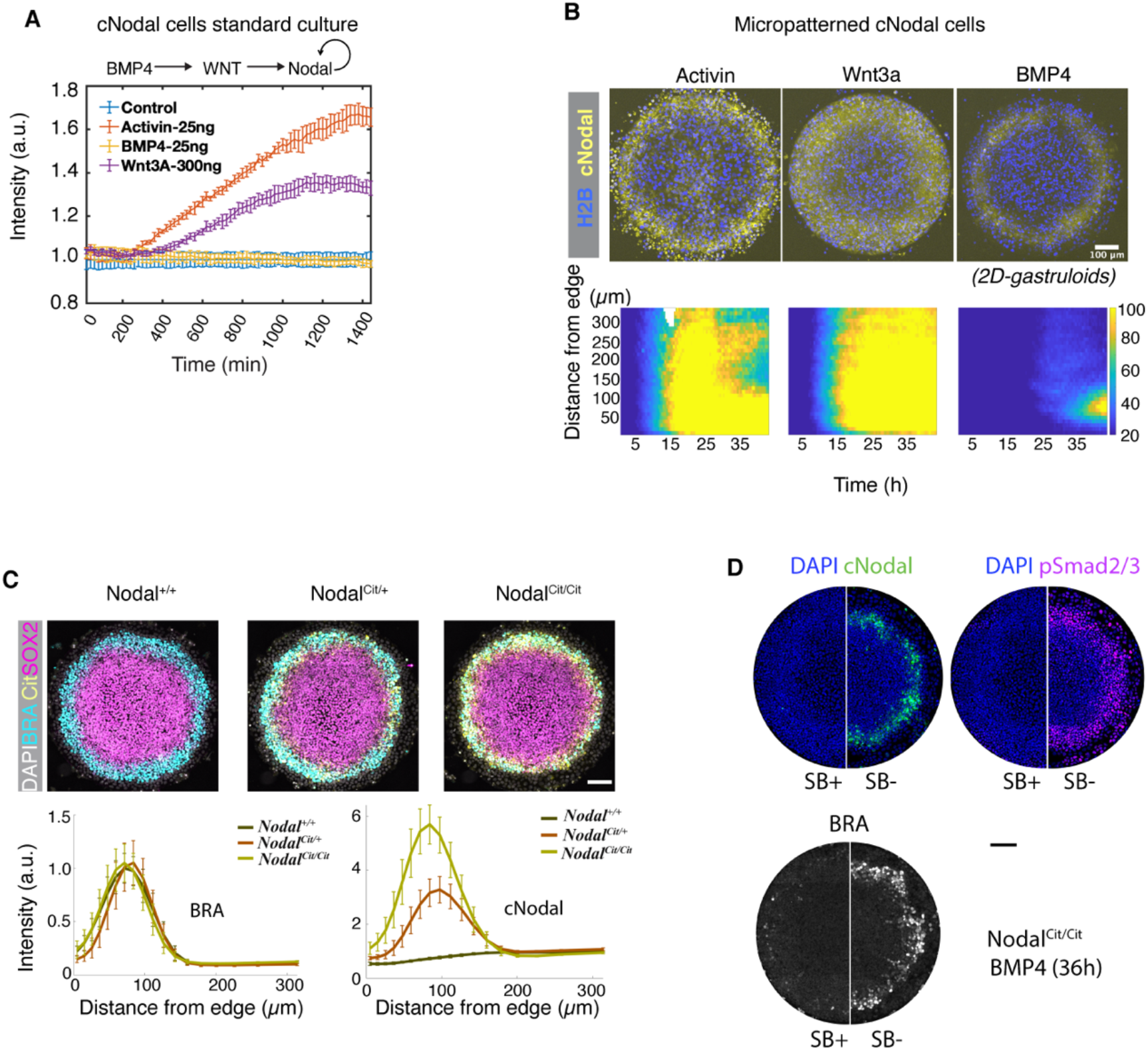
Visualization of a fully functional endogenous Nodal protein in hESCs and gastruloids. (A) Quantification of cNodal expression in response to addition of Activin, Wnt3a, or BMP4. Normalized mean intensity of cNodal from time-lapse imaging over 24h with a 20-minute interval is quantified (n=5). (B) Activin, Wnt3a or BMP4 treated micropatterned cNodal hESCs with CFP-H2B nuclear marker. (bottom) Mean intensity of cNodal from time-lapse imaging over 43h with 30 minute interval is quantified as function of time and distance from colony edge. (C) Quantification of BRA and cNodal levels in gastruloids made with the indicated genotypes (n=10). Brachyury (BRA) represents mesodermal fate. SOX2 represents ectodermal fate. Cit represents anti-mCitrine (GFP antibody). (D) SB-431542 a type I ALK receptors inhibitor was used to inhibit Nodal signaling in homozygous cells and the expression of cNodal, pSmad2/3 and BRA were compared (representative of n=6 for each condition). Scale bar, 100 µm. Maximum intensity projection images are shown in all figures. Data in (C,D) is at 40h differentiation.

We utilized cNodal hESCs to make gastruloids. We found that cNodal heterozygotes and homozygotes give rise to normal fate patterns indistinguishable from wild type (WT) patterns (Fig 1C), while gastruloids made with Nodal knockout hESCs lack the mesodermal layer (*20*) (Fig. S4), consistent with the phenotype of Nodal^-/-^ mice (*4*). Moreover, Nodal^cit/cit^ gastruloids show normal patterns of activated Smad2/3 (Fig. 1D, Fig. S5), while we have previously observed significantly lower levels in Nodal^-/-^ cells (*20*). Together, these results demonstrate that the cNodal fusion is fully functional.

### Nodal and Lefty propagate by a relay mechanism

Next, we surveyed the ranges of Nodal and Lefty proteins through juxtaposition of cNodal cells (senders) with Lefty1/2 null cells (receivers) (Fig. 2A-C, Fig. S6-7) (*21*). Activin pretreated senders produce both Lefty and cNodal, while receiver cells do not produce either. Therefore, the spatial range of the two proteins can be obtained via measuring fluorescence intensity in the receivers as a function of the distance from the senders. We failed to observe Nodal protein beyond the receiver cells immediately adjacent to the senders (Fig. 2B, C and Fig. S7C), while Lefty traveled over 6-8 cell tiers from the source and maintained stable gradients over time (Fig. 2C and Fig. S7C).

**Figure 2.**
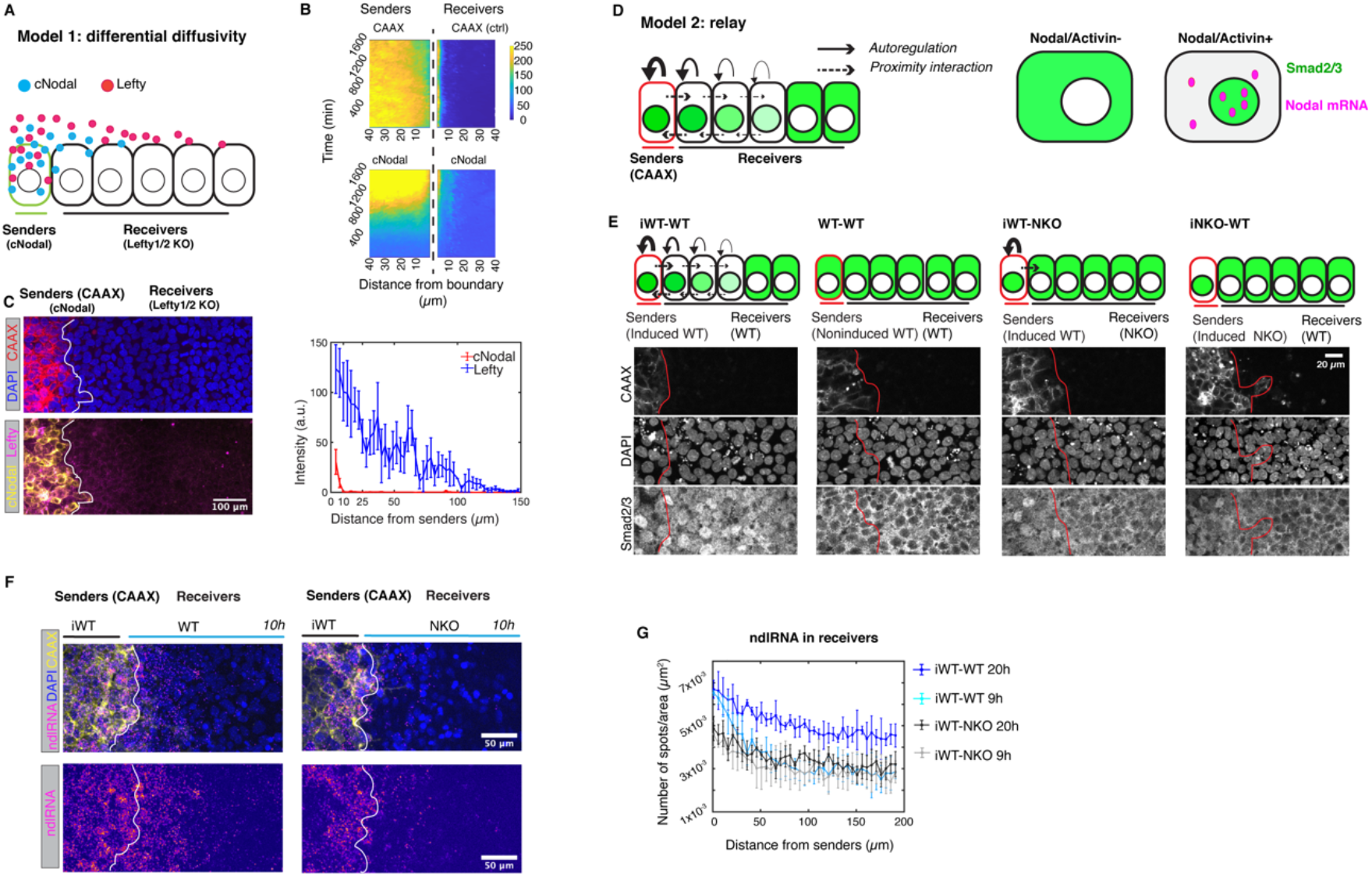
Nodal propagates by a relay mechanism. (A) Schematic of protein differential diffusivity model. Senders (green outline) represent cells that produce cNodal (solid blue circle) and Lefty1/2 (solid magenta circle) upon Activin induction. Receivers represent Lefty1/2 KO cells that cannot make either Lefty or cNodal proteins. (B-C) Experimental test of the differential diffusivity model. Sender cells are labeled by cell membrane localized mCherry-CAAX. (B) Live-cell imaging over 27h with a 20-minute interval. mCherry-CAAX and cNodal levels in senders or receivers are quantified as function of distance from border. mCherry-CAAX senders as a non-diffusive control. (C) Anti-GFP that recognizes mCitrine and anti-Lefty antibodies (Fig. S6, 7) were used to measure protein ranges. Quantitative analysis of staining images is shown. Mean intensity of cNodal (anti-GFP) and Lefty at 20h is quantified as a function of distance from border (n=4). Scale bar, 100 µm. Images are 20h post juxtaposition. (D) Schematic of relay model. Senders (red outline) represent induced cells that express Nodal. Solid line with arrowhead, represents Nodal positive autoregulation. Dashed lines with arrowheads represent juxtacrine induction by Nodal. Nodal/Activin signaling transducers Smad2/3 (green) are located in cytoplasm in the absence of ligands. In the presence of Nodal or Activin, Smad2/3 translocate to nucleus, meanwhile Nodal transcription is upregulated. Nodal mRNA is shown as magenta solid circle. (E-G) Experimental test of the relay model. (E) Staining of Smad2/3 to measure Nodal signaling activity in juxtaposition experiments with cells with the indicated genotypes. Signal is nuclear when the pathway is activated. Sender cells in all conditions were labelled by mCherry-CAAX. Images in each case taken after 20h juxtaposition n=4. (F-G) Experiments performed as in (E) and Nodal mRNA (ndlRNA) was measured. Nodal RNA was quantified as a function of distance from border (n=4 for each condition) (F) Representative images of 10h post-juxtaposition is shown, scale, 100 µm. (G) Nodal RNA was measured in a separate experiment with 9h and 20h post-juxtaposition. iWT means induced wild type cells, iNKO means induced Nodal KO cells.

The failure of Nodal protein to spread from source cells seemingly contradicts our previous observations that Nodal signaling activity can propagate from the edge to the center of gastruloids, even in the absence of upstream Wnt signaling (*20, 22*). This discrepancy is potentially consistent with the idea that signaling spreads primarily through a relay mechanism in which Nodal activity induces transcription of Nodal ligand in immediately adjacent cells (Fig. 2D).

To test the relay hypothesis (Fig. 2D), juxtaposition experiments with various combinations of sender and receiver hESCs were carried out (Fig. 2E). In particular, we induced Nodal expression in sender cells by treating them with Activin and then placed them adjacent to either wildtype or Nodal^-/-^ cells. As controls, we also used non-induced cells or induced Nodal^-/-^ cells (NKO) as senders. Dramatically elevated levels of Smad2/3 nuclear accumulation were only observed when both the sender and receiver cells had a functional Nodal protein, either Nodal^**+/+**^ or Nodal^Cit/Cit^ (Figure 2E). When Nodal^-/-^ cells were used as receivers, the signal was not transmitted beyond the receiving cells which touch the senders, suggesting that a relay involving induction of Nodal ligand in the receivers is required. Non-induced and Nodal^-/-^ cells also failed to induce a response showing that the response is specific to Nodal from the senders. We also measured the induction of Nodal and Lefty transcripts in a similar assay (Fig. 2F, Fig. S7E-F). Induced WT senders were juxtaposed with non-induced WT cells or with Nodal^**-/-**^ cells for 9-20h hours. In the former case (WT - WT), we observed gradients of Nodal RNA, Lefty RNA and nuclear Smad2/3, which peaked at the border and declined with distance from the senders. However, when Nodal^-/-^ cells were used as receivers, we did not observe elevated signaling activity or increased Nodal or Lefty mRNA production (Fig. 2F-G, Fig. S7E-F). Interestingly, lower levels of Nodal RNA and Lefty RNA were observed in the sender cells when juxtaposed with Nodal^**-/-**^ cells, and by 20 hours, induction of these Nodal targets was largely absent in the senders juxtaposed to Nodal^**-/-**^ cells but maintained in senders juxtaposed to wildtype cells. These results indicate that propagation of signal is bidirectional and that signaling from the receivers is required to maintain Nodal expression in the senders. Together, these results support that a transcriptional relay is required for the propagation of Nodal signaling in space.

We then sought to test whether this hypothesis stands true in micropatterned human gastruloids. In this model, self-organized patterns are formed with extraembryonic cells at the colony edge, ectoderm at the center, and rings of mesendoderm in between. We first assessed Nodal and Lefty transcription in space and time (Fig. 3A). The results showed that peak expression of both Nodal and Lefty mRNA was first detected at the colony edge, then both expanded towards the colony center as development of the gastruloids proceeded. Although both moved inwards, Nodal mRNA expanded to cover a broader range while remaining high at its point of initiation, consistent with measurements of Smad2 signaling activity (Fig. 3C, Fig, 4A, Fig. S8 and (*20, 22*)). In contrast, Lefty RNA, a direct Smad2-dependent Nodal signaling target, shifted rapidly inward so that it was only expressed transiently at each spatial position, and the peak active position moved continually inward. These results show that Nodal activity spreads inwards to cover a domain of increasing size driven by expression of Nodal mRNA, while Lefty mRNA is only expressed at the front of this spreading domain in the cells which have most recently activated Nodal signaling. These data suggest that Lefty expression is adaptive, increasing upon initial exposure to Nodal and then returning to low levels despite continued Nodal protein expression and signaling activity (Fig. S13).

**Figure 3.**
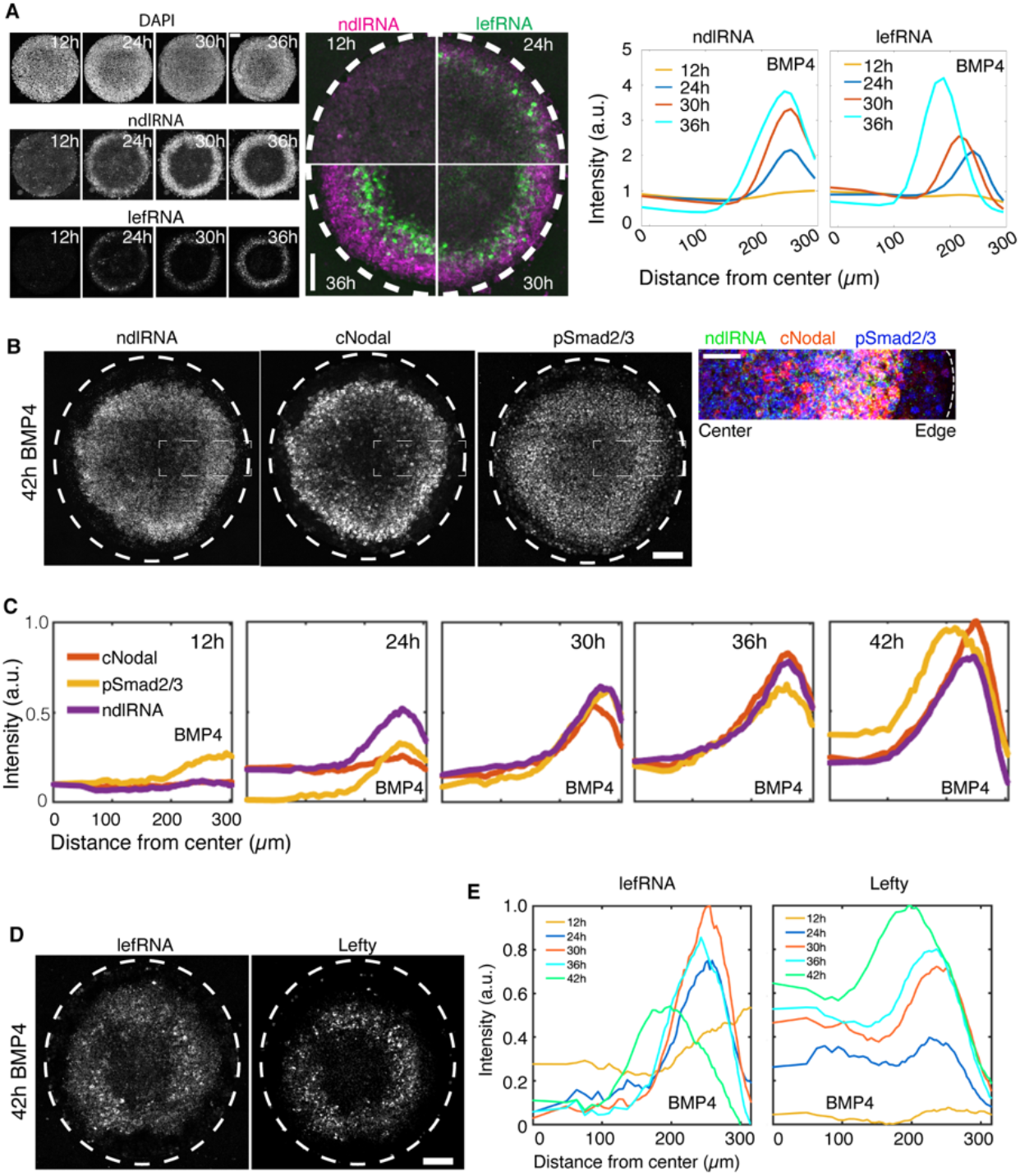
Nodal and Lefty spread via a relay mechanism in gastruloids. (A) smFISH results showing mRNA for Nodal and Lefty1/2 at 12h, 24h, 30h, 36h post-BMP4 treatment. (right) Quantitative analysis of the smFISH mean intensity as a function of distance from colony center at indicated time (n=10), scale bar, 100 µm. (B) Images of gastruloids at 42h, showing distribution of RNA and protein of Nodal and pSmad2/3, scale bar, 100 µm. The rectangular box indicates the enlarged area in the right panel, scale bar 50 µm. (C) Simultaneous quantification of Nodal RNA and protein and pSmad2/3. Normalized mean intensity is quantified over time as function of distance from colony center (n=6). (D) Images of gastruloids at 42h, showing Lefty mRNA and protein. (E) Quantification of Lefty RNA and protein over time, mean intensity as function of distance from colony center (n=6). Scale bar, 100 µm.

Next, by simultaneously tracing Nodal mRNA, Nodal protein and phosphorylated Smad2/3 (pSmad2/3) (Fig. 3B, C), we found that, in line with the relay model, Nodal protein and activity were only found in the region of Nodal mRNA production, suggesting essentially no transcriptionally-independent spread (Fig. 3C, Fig. S8, 9). Interestingly, in contrast to its broader range in the juxtaposition assay, Lefty protein moved as a traveling wave, closely mirroring the mRNA wave (Fig. 3D, Fig. S9), suggesting that in this context, Lefty remains close to its source and moves inwards primarily through a shift in the region of transcription of Lefty mRNA. The difference between these assays may reflect the different time scales involved. In the juxtaposition assay, the Lefty gradient forms over 20 hours (Fig. S7C), while in the gastruloid model, transient expression at each spatial position might not allow for spread of Lefty. Taken together, the results indicate that shifting patterns of Nodal and Lefty reflect a transcriptional relay, while extracellular movement of these proteins plays little to no role in patterning micropatterned gastruloids.

### Lefty restricts Nodal in space and time

To provide clues on how Nodal and Lefty self-organization patterns human gastruloids, we created gastruloids with WT hESCs or Lefty1/2 compound knockout (Lefty1^-/-^, Lefty2^-/-^) hESCs (Fig. 4). The results show that Lefty protein ablation led to upregulated Nodal production at 12h with high levels peaking at the colony edge and extending in a shallow gradient towards the center from 24h to 42h. In contrast in WT gastruloids, Nodal transcripts did not emerge at elevated levels until 24h and then peaked at colony edge with fronts gradually propagating inwards, (24-42h). In line with Nodal expression patterns, comparing to WT gastruloids, pSmad2/3 levels were detected earlier and in a broader range extending to the colony center of mutant gastruloids (Fig. 4A). In addition, analysis of Lefty transcripts shows that sustained pSmad2/3 levels did not stimulate Lefty transcription (Fig. S13), which supports the hypothesis discussed above of adaptive Lefty expression.

**Figure 4.**
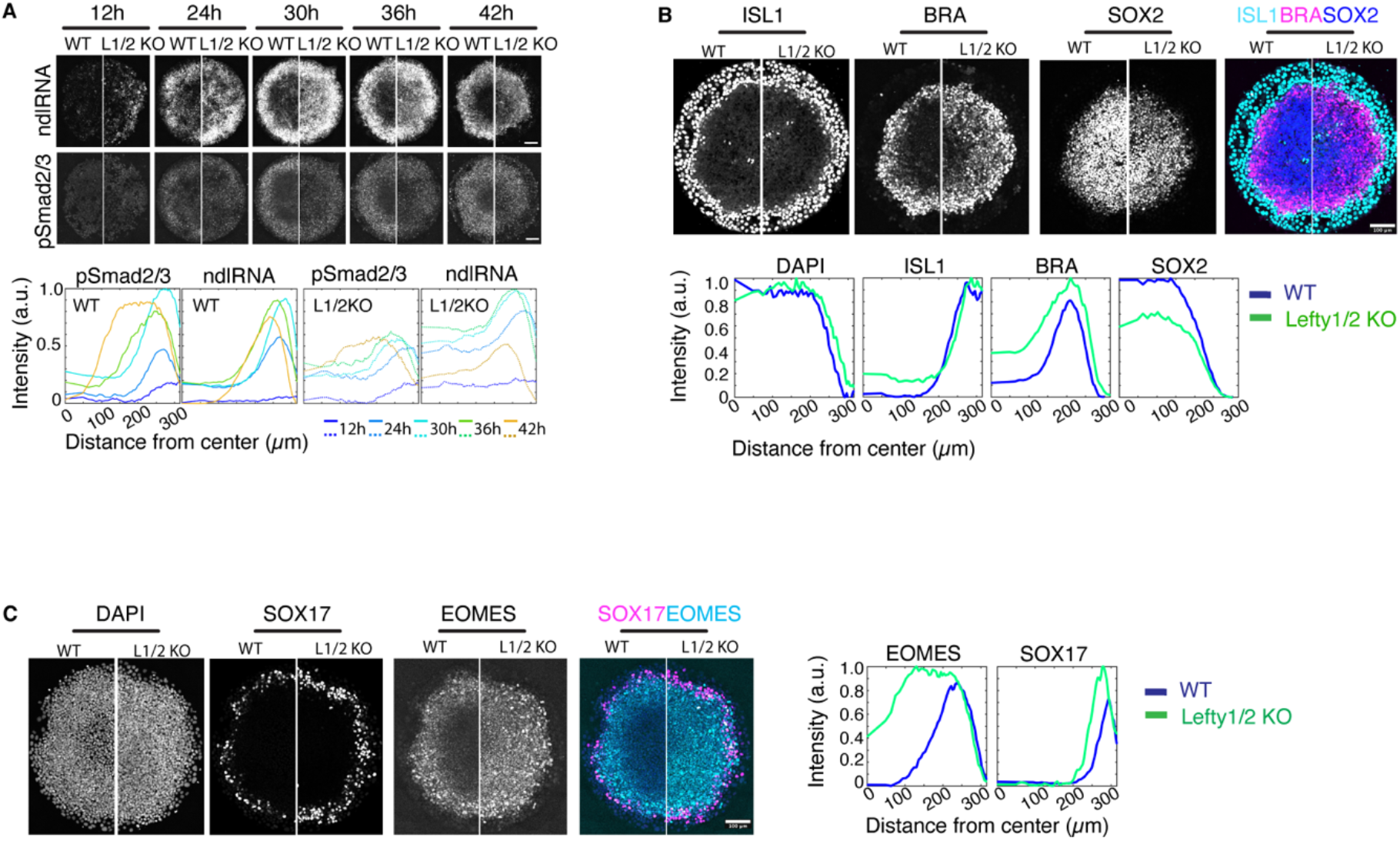
Lefty is needed to slow spread of Nodal and separate mesoderm and ectoderm. (A) Comparison of Nodal mRNA expression and pSmad2/3 over time in gastruloids created with WT and Lefty1/2 compound KO hESCs. Representative images at all time points are shown. (B) Immunofluorescence for ISL1, extraembryonic marker. BRA, mesodermal marker, and SOX2, ectodermal marker. (C) Immunofluorescence for SOX17, definitive endodermal marker, and EOMES, mesendodermal marker. In all cases, normalized mean intensity is quantified as a function of distance from colony center (n=6 for each condition). Images in (B,C) acquired after 40h of BMP4 treatment. Scale bar, 100 µm in all panels.

Previous studies have shown that a sequence of signaling activities through the BMP, Wnt, and Nodal pathways are responsible for patterning the gastruloids (*20, 23*). To verify whether or not the altered Nodal signaling is directly caused by Lefty depletion or might also involve changes in upstream signaling, we assessed BMP4 and Wnt signaling in Lefty knockout gastruloids. The results show that BMP4 and Wnt signaling largely remain qualitatively unchanged (Fig. S12). Therefore, altered Nodal signaling is not likely caused by perturbation to upstream pathways. These results together suggest that Lefty is required to restrict Nodal signaling in space and time.

### Lefty inhibition to Nodal is required for Mesodermal patterning

We then further investigated the role of Nodal-Lefty self-organization in fate patterning by surveying germ layer differentiation in control and Lefty KO gastruloids (Fig. 4B, C). The data show that Lefty depletion results in expanded mesodermal (BRA+) fates at the expense of ectodermal cells (SOX2^+^). Interestingly, the SOX17^+^ boundary is unchanged between WT and Lefty1/2 KO. Mesendoderm differentiation was also detected earlier and more broadly in Lefty1/2 mutant gastruloids with BRA expression observed in the center where ectoderm usually forms at 24h in Lefty1/2 mutant gastruloids, but not in WT gastruloids (Fig. S10-11), suggesting that the early upregulation of Nodal signaling in this region drives rapid differentiation.

Here we have observed endogenous Nodal and Lefty in hESCs and during gastruloid patterning. Visualization of protein together with the encoding mRNA allowed us to determine the range of action of these molecules. Nodal does not travel more than one cell diameter in any context we examined, and, Lefty protein expression, while capable of forming a stable gradient, closely mirrors that of Lefty mRNA during gastruloid patterning. Nodal moves by a transcriptional relay mechanism, and patterns of transcription largely account for both Nodal and Lefty spreading. This mechanism is likely different from the mode of action of Nodal in Zebrafish where it has been shown that Nodal can spread at a long distance without requiring signaling in target cells, and its range is typically limited by coreceptor expression (*24*). Differences in effective Lefty range during patterning may also reflect different developmental strategies: in Zebrafish long range Lefty is needed for scaling fate patterning to embryo size (*25*), while the patterns in human gastruloids do not scale (*16, 26*). Taken together, our study defines a new mechanism of patterning by an activator-inhibitor pair that operates in specifying cell fates in human gastruloids.

