## Supplementary information for "Nodal is a short-range morphogen with activity that spreads through a relay mechanism in human gastruloids"

### Materials and Methods

#### Cell line

##### Human embryonic stem cell (hESC) line

ESI-017 hESC line used in this study was purchased from ESIBIO. Karyotype and genomic integrity check were done by the manufacturer. Pluripotency (OCT4+, SOX2+, NANOG+) was regularly monitored prior to and during the study. Nodal knockout cells were previously generated and characterized in this lab (*1*). All cell lines were tested regularly for mycoplasma and found negative.

#### Cell culture

hESCs were maintained in serum-free, mTeSR1 (STEMCELL Technologies) medium, in Matrigel (Corning; 1:200 in DMEMF12) coated culture dishes, at 37°C, 5% carbon dioxide (CO2). Essential 6 medium (E6) (Gibco) was used for TGF-ß ligand-free conditions (Figure S11). All experiments were performed with cells not exceeding passage number 60. Dispase (Fisher Scientific) for gentle dissociation was used for routine maintenance. Accutase (Corning) was used to prepare single cell suspensions. Rock-inhibitor Y27672 (10 µM; StemCell Technologies) was used to support single cell viability.

Antibiotics were used for selection. In particular, 100 µg/ml G418 or 5 µg/ml Blasticidin was used to select cells with a modified Nodal locus; 1 µg/ml Puromycin was used to enrich for cells transiently transfected with px459 plasmids; 5 µg/ml Puromycin was used for selecting the mCherry-CAAX cell membrane marker. 5 µg/ml Blasticidin was used for the H2BCFP nuclear marker. Inducible expression experiments were performed in mTeSR medium supplemented with the indicated factors for the indicated times.

**Plasmids**

Plasmids were constructed for this study as follows.

1) ePiggyBac transposable element vectors for stable integration of genes of interest (*2*). An ePiggyBac master vector based on the pBSSK backbone, harboring transposon-specific inverted terminal repeat sequences (ITRs) was modified to deliver: nuclear marker H2BCFP (Plasmid AW-P28), data shown in Figure 1B; doxycycline inducible cell membrane marker mCherry-CAAX (Plasmid AW-P224), data shown in Figure 1D and 2C.

2) CRISPR/Cas9 vectors for genome-editing. To generate insertions (cNodal) or nonsense mutations (Lefty 1/2), a Cas9 and sgRNA co-expression vector, either px459 (Addgene) or px330 (Addgene), was modified to target the locus of interest: Nodal mature domain targeting plasmid AW-P188, derived from px330, sgRNA spacer sequence 5'-TGTCTGGCAAGTGATGTCGA-3'; Lefty 1 exon1 targeting plasmid AW-P219, derived from px459, sgRNA spacer sequence 5'-GCAGCACCATGCAGCCCCTG-3'; Lefty 2 exon1 targeting plasmid AW-P220, derived from px459, sgRNA spacer sequence 5'- GCAGCACCATGTGGCCCCTG-3'. All sgRNA sequences used in this study were designed using the Benchling built-in function Design CRISPR Guide. Off-target score was evaluated using the tool found at crispr.mit.edu from (*3*). On-target score was evaluated according to the optimized score from (*4*), only available for SpCas9.

3) Plasmids for homology directed DNA repair. Two donor vectors were created for biallelic Nodal locus insertion. The configuration of the two vectors is shown in Figure S1A. In the FloxP-neomycin resistant vector, 800bp of Nodal pro-domain or Nodal mature domain sequence was used as right or left homology arm, respectively, flanking the FloxP-neomycin expression cassette

and mCitrine. In the FloxP-blasticidin-2A-RFP vector, 800bp of Nodal pro-domain was used as right-arm, 450bp of Nodal mature domain was used as left-arm. Specifically, refer to plasmid AW-P216 (Neo) and AW-P218 (Bsd-RFP). Cre expression plasmid (Cre Shine, Addgene) was used for excision of FloxP fragments after antibiotic-resistance selection.

**DNA transfection and stable cell line establishment**

DNA transfection was performed with the 4D-Nucleofector (Lonza) system according to the manufacturer’s protocol using the P3 Primary Cell 4D-Nucleofector Kit (Lonza). For ePiggyBac transposase mediated insertion, antibiotic selection started on day 2 (two days after nucleofection), and lasted at least for 7 days. For CRISPR/Cas9 mediated knockout, px459 plasmid transfectants was selected on day 1, for one day. For CRISPR/Cas9 mediated insertion, donor DNA was linearized and concentrated before nucleofection. Selection started on day 3, and lasted for about 7-10 days, until no cell death was observed and healthy colonies were established.

For ePiggyBac transposase mediated insertion, cells were pooled after selection. Corresponding antibiotics were not supplemented in medium for routine culture, until 2-3 days prior to an experiment. For CRISPR/Cas9 mediated knockout, single clones were handpicked and amplified. Sub-cloning was conducted as needed. Sanger sequencing was performed to screen retrieved colonies. Promising knockout candidates were further confirmed by sequencing individual alleles with TOPO-cloning (Invitrogen). Only confirmed knockout mono-clones were used for further experiments. For CRISPR/Cas9 mediated insertion of mCitrine to Nodal locus, the insertion procedure was done for twice to create the homozygous knock-in line. The modified locus is not recognized by the sgRNA so the first knock-in allele was not edited in producing the second. Single-cell Fluorescence-activated cell sorting (FACS) was conducted to isolate desired mono-clones, with correct genomic modification. After establishment, stable lines were checked for pluripotency markers, i.e., OCT4, SOX2 and NANOG expression, and found indistinguishable from WT ESI-017.

**Single-cell FACS**

We found that transiently expressed Cre-mediated excision of FloxP fragments inserted in Nodal locus in hESCs was extremely inefficient. Therefore, desired clones with FloxP excised could only be isolated from post-Cre mixture by single-cell FACS. A SH800S Cell Sorter (Sony) was used to sort single cells into individual wells of 96-well plates. Potential heterozygotes were treated with 10 ng/ml Activin in order to increase cNodal expression, so that mCitrine could be used as marker for FACS. Potential homozygotes were also pre-treated with Activin and sorted as mCitrine-positive/RFP-negative. Sorted single cells were kept in mTeSR medium supplemented with 1x CloneR (STEMCELL Technologies), to support viability and genomic integrity. Mono-clones were established and characterized by genotyping PCR and Sanger sequencing (shown in Figure S1).

**Polymerase chain reaction (PCR)**

PCR for DNA fragment sub-cloning for plasmid DNA construction was done using OneTaq Hot Start (NEB), Phusion Hot Start (NEB) or Q5 (Fisher Scientific) DNA polymerase, following the manufacturer's instructions with optimized annealing temperature. Optimized conditions were used for Nodal modified cells genotyping PCR (*5*).

**Micropatterned hESCs-based gastruloids**

We used CYTOOplates 96 DC-S-A glass-bottom 96-well microplates with circular micropattern (diameter, 700 µm) for creation of gastruloids. On the day of seeding cells, micropattern was coated with 200 µl 5 µg/ml laminin-521 (LN521, Biolamina) in DPBS (with calcium and magnesium, Lonza) for 2 h at 37°C. Unattached laminin was washed out with 200 µl of DPBS, repeated 4 times.

hESCs were passaged using Dispase 2-3 days prior to micropatterned cell seeding and maintained in mTeSR medium. By the day of seeding, cells reached 50-70% confluency, with the majority of the colonies 500 – 1000 µm in diameter. Single cell suspension was prepared by using Accutase, and cells were counted using a hemocytometer. Unless specifically indicated, about 150,000 cells/150 µl mTeSR were placed into each well of the laminin coated micropattern plate, and incubated at 37°C for 45 minutes to allow the cells attached to micropattern surface. Then, unattached cells were washed away with PBS twice, before adding induction medium. The induction medium is mTeSR supplemented with 50 ng/ml BMP4, unless otherwise specified. For detailed protocol, refer to (*6*).

**Juxtaposition analysis**

For cNodal and Lefty protein diffusivity analysis, homozygous cNodal hESCs or Lefty 1 and Lefty 2 (Lefty1/2) compound knockout hESCs (for background assessment) were used as senders. Lefty1/2 compound knockout hESCs were used as receivers. For temporal analysis experiments, sender cells were dissociated with Accutase and 100,000 cells were resuspended in 50 µl mTeSR supplemented with Rock-inhibitor Y27672 (10 µM) and seeded in one well of 2 well silicone insert with 0.22 $cm^2$ growth area (ibidi), in a Matrigel coated µ-Slide 8 Well (ibidi), and then incubated at 37°C for 1-2 hours to allow the cells attached to culture dish surface. Unattached cells were washed out with PBS twice. Sender cells were then induced with 50 ng/ml Activin in mTeSR for 20h. The following day, Lefty1/2 compound knockout cells were dissociated with Accutase and 200,000 cells were resuspended in 100 µl mTeSR supplemented with Rock-inhibitor Y27672 (10 µM), and seeded in the same well, juxtaposed with sender cells. Unattached receiver cells were washed out after 30 minutes incubation with PBS. Then, 150 µl mTeSR supplemented with or without 50 ng/ml Activin was added into the well. Cells were fixed at indicated time points for further assessments of cNodal and Lefty protein.

For positive autoregulation-dependent relay analysis. WT hESCs or homozygous cNodal hESCs were used as positive sender cells, Nodal knockout hESCs were used as negative sender cells. WT hESCs or Nodal knockout hESCs were used as receiver cells. For assays under stationary conditions, sender cells were seeded in silicone insert and kept in mTeSR supplemented with 10 ng/ml Activin for 16-20 hours. Prior to seeding of receiver cells, sender cells were washed with PBS for two times, and then with mTeSR or E6 for one time. Specific receiver cells were seeded juxtaposed with sender cells. Cells were kept in media alone (mTeSR or E6), and then were fixed at indicated time for further analysis of Smad2/3, Nodal transcripts or Lefty transcripts. Of note, to minimize basal levels of Nodal expression caused by TGF-ß in mTeSR, the Nodal FISH experiment shown in Figure 2F was done with sender and receiver cells kept in E6 medium. In addition, 10 μM SB431542 was added into the medium at 1h prior to dissociation of receiver cells. In that case, the receiver cells were washed two times with E6 after accutase digestion to remove SB431542.

Heterogeneous or decreased Nodal expression or signaling levels were occasionally observed in the sender cell colony center after long incubation times (after 24-48h hours), which might be caused by high cell density (*7*). To avoid these effects complicating the analysis, freshly seeded sender cells were induced with Activin for 16-20h or relatively low-density cells at the colony edge of sender cell colonies were imaged as signal sources.

**Single molecule fluorescence in situ hybridization (smFISH)**

Cells were fixed in 4% paraformaldehyde for 15 minutes. smFISH were performed following manufacturer's instructions (ACD Bio, RNAscope fluorescent multiplex assay), with indicated probes (Table S1). Cells were placed in PBS before imaging.

**Immunofluorescence staining**

Cells were fixed in 4% paraformaldehyde for 15 minutes at room temperature. Fixed samples were then washed with PBS twice before permeabilization and blocking with 3% Donkey serum in PBST (1xPBS with 0.1% Triton X-100) for 30-60 minutes at room temperature. For pSmad2/3 staining, cells were incubated in PBS with 1% SDS for 30 minutes at 37°C following blocking.

Primary antibodies were diluted in blocking buffer as indicated (Table S1). Cells were incubated with diluted primary antibody solution at room temperature for 1 hour or at 4°C overnight, washed three times with PBST for 30 minutes each time, and then incubated with 1:500 diluted secondary antibodies (Table S1) solution supplemented with DAPI at room temperature for 1 hour, and then washed with PBST 3 times. Cells were placed in PBS before imaging.

For dual smFISH/immunofluorescent labelling, the smFISH procedure was performed first and then cells were washed with PBS and incubated with primary antibodies. The remainder of the immunofluorescence procedure specified above was then followed.

**Imaging**

*Live cell imaging*. Cells seeded in μ-Slide 18 well or 8 well (ibidi) chamber slides or CYTOO micropattern 96-well plates were imaged on Olympus/Andor spinning disk confocal microscope equipped with environmental chamber, with a 20x, NA 0.75 objective or 40x, NA 1.25 silicon oil objective. During imaging, temperature (37°C), humidity (~50%), and $CO_2$ (5%) were controlled. 4 positions of each conditions were selected for imaging.

Fixed cell imaging. Fixed micropatterned cells were imaged on Olympus/Andor spinning disk confocal microscope, using 20x, NA 0.75 objective. 6 to 10 colonies were imaged for each condition and 4 positions were imaged for each colony so that the entire colony could be covered and reconstituted; Olympus IX83 inverted epifluorescence microscope and Olympus FV1200 laser scanning confocal microscope (LSM) with 20x, NA 0.75 objective were also used for fixed micropattern imaging. Fixed cells in chamber slides were imaged on spinning disk confocal microscope with a 20x or 40x, NA 1.25 silicon oil objective or epifluorescence microscope with 20x objective. 4 positions were imaged for each condition.

**Image analysis**

Imaging experiments were repeated at least twice with consistent results. Images obtained from various experimental conditions were processed using Fiji (*8*) to set lookup tables for visualization.

Filters (Remover outliers) were applied to some raw images to remove non-specific bright puncta in lookup tables. Ilastik (*9*) was used for image segmentation. Analysis of micropattern images and live-cell imaging in standard culture analysis was performed using custom MATLAB code, available at https://github.com/warmflashlab/ImageProcessing. For images obtained from juxtaposition experiments custom MATLAB code, available at https://github.com/anemashkalo/lefty-nodal-.git. Briefly, images were first segmented via Ilastik to detect all nuclei (via DAPI channel). The channel representing one of the two cell types (usually the sender cells) was also segmented. Based on this, binary image masks were created to contain either only producing or only receiving cell areas. Images were background-subtracted and protein expression was quantified in receiving cells as a function of distance from producing cells in several-micron increment slices away from the border between cell types. Since markers were quantified in both nuclear and cytoplasmic space, DAPI normalization was not used in these cases. Distance from the border was determined using the MATLAB function *bwdist* applied to producing cells' mask. smFISH images were quantified in similar way, instead of separating individual molecules, average pixel intensity was used. To quantify smFISH in standard culture (Fig. 2F) mean pixel intensity was quantified from masks generated via cell nuclei segmentation and dilation, in order to account for cytoplasmic mRNA molecules. Error bars represent SEM over multiple images at each time point. Kymographs of micropatterned images (in Fig. 1B) were obtained as averages over three circular colonies containing four pie-segments each, i.e averaged over 12 pie segments for each treatment condition.

**Statistical analysis**

Multiple images of micropatterns or juxtaposition cultures or FISH experiments contributed to each experimental condition. In quantification of protein / mRNA expression as a function of distance from cell types border, error bars represent standard error of the mean (SEM) over individual images for each experimental condition. In quantification of cNodal protein expression over time shown in Figure 1A, C, error bars represent standard deviation (SD).

**Table 1 Key reagent or resource**

| Reagent | Dilution / concentration | Source | Identifier |
|---|---|---|---|

| | | | |
|---|---|---|---|
| Mouse anti-GFP | 1:200 | Abcam | Cat# ab1218 |
| Mouse anti-Smad2/3 | 1:200 | BD Biosciences | Cat# 610843 |
| Mouse anti-ISL1 | 1:75 | DSHB | Cat# 39.4D5 |
| Mouse anti-Oct3/4 | 1:400 | BD Biosciences | Cat# 611203 |
| Rabbit anti Eomes | 1:200 | Abcam | Cat# ab23345 |
| Rabbit anti-pSmad1/5/8 | 1:100 | Cell Signaling | Cat# 13820 |
| Rabbit anti-pSmad2/3 | 1:100 | Cell Signaling | Cat# 8828S |
| Rabbit anti-Smad2 | 1:200 | Cell Signaling | Cat# 5339S |
| Rabbit anti-SOX2 | 1:200 | Cell Signaling | Cat# 5024S |
| Goat anti-BRA | 1:300 | R&D Systems | Cat# AF2085 |
| Goat anti-HAND1 | 1:200 | Cell Signaling | Cat# AF3168 |
| Goat anti-Nanog | 1:200 | R&D Systems | Cat# AF1997 |
| Goat anti-Lefty | 1:200 | R&D Systems | Cat# AF746 |
| Goat anti-SOX17 | 1:200 | R&D Systems | Cat# AF1924 |
| Activin A | 10-100 ng/ml | R&D Systems | Cat# 338-AC |
| BMP4 | 50 ng/ml | Fisher Scientific | Cat# 314BP050 |
| Wnt3a | 200-300 ng/ml | R&D Systems | Cat# 5036-WN |
| SB431542 | 10 µM | Stemgent | Cat# 04-0010-05 |
| Nodal RNAscope probe | 1x | ACD Bio | Cat# 416541-C3 |
| Lefty RNAscope probe | 1x | ACD Bio | Cat# 415651 |
| Axin2 RNAscope probe | 1x | ACD Bio | Cat# 400241-C3 |
| Lefty1 siRNA | N/A | Dharmacon | Cat# L-013114-00-0005 |
| Lefty2 siRNA | N/A | Horizon Discovery | L-017473-00-0005 |

**Table 2 Primers used for genomic DNA PCR**

| **Target gene** | **Forward** | **Reverse** |
|---|---|---|
| Nodal | 5' GCATGGTTTTGGAGGTGACCAG 3' | 5' ACATGCCTGGTACCTAGCACAG 3' |

| | | |
|---|---|---|
| Nodal | 5' ATGCTCTACTCCAACCTCTCG 3' | 5' ATGTATGCATGGTTGGTC 3' |
| Lefty1 | 5' CAGCTGGGGGAACTGAGCCTCT 3' | 5' TCAGCCTCCCACAGACCTCTGC 3' |
| Lefty1 | 5' AGACCACTGCCCTCCAGTG 3' | 5' AACACCAGCAGGTGTGTGC 3' |
| Lefty2 | 5' ATTGTCAAGCTGCTGACAGC 3' | 5' AGCAGGACTACATACTGGGC 3' |

Supplementary figures

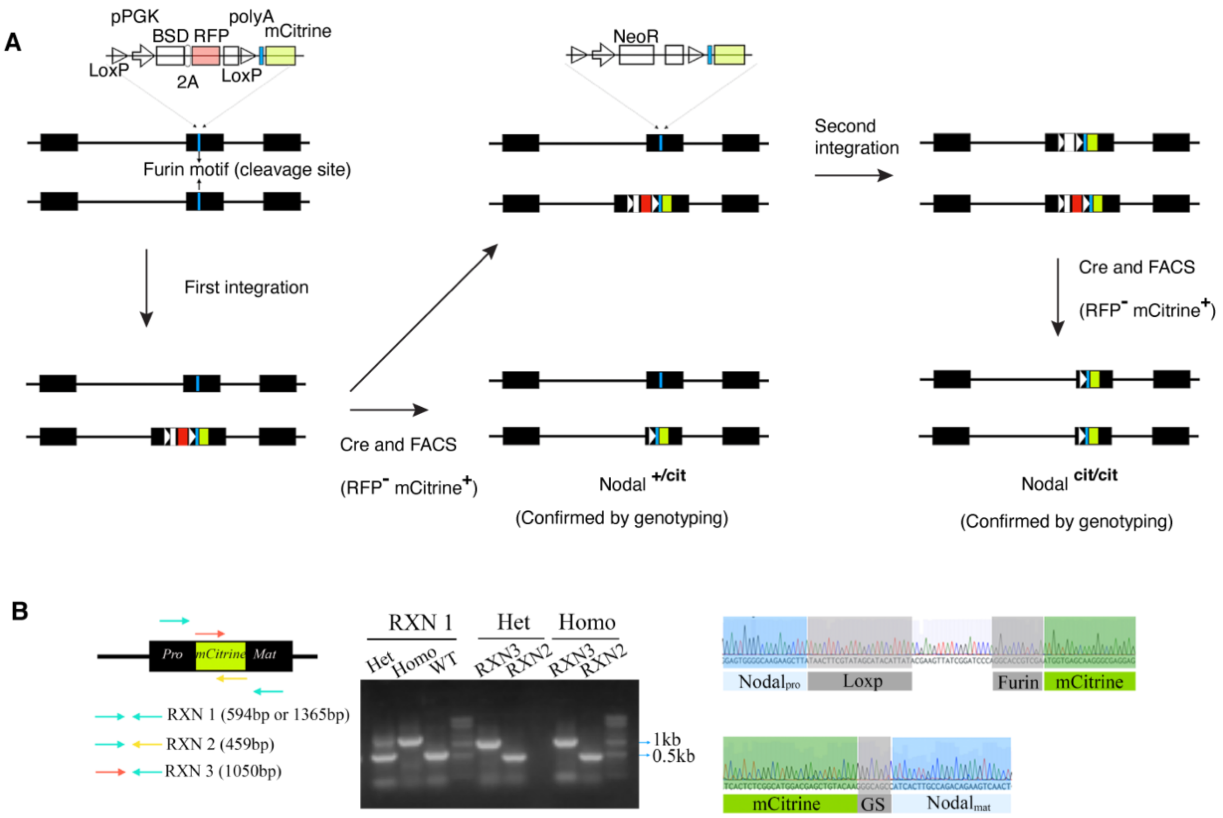


**Supplementary figure 1. Insertion of mCitrine to targeted Nodal locus.** (A) Schematic of CRISPR/Cas9 mediated double-strand DNA break (DSB) repair by homologous recombination. The cleavage site (blue vertical bar) is located within Exon2 of the *NODAL* gene, specifically, within the last codon of the 'RHRR' Furin consensus motif. Linearized donor DNA with 400-800 bp homology arms was provided for DSB repair. The donor DNA is composed of the fluorescent protein and loxP sites flanking an antibiotic resistance gene expression cassette, i.e., blasticidin S deaminase (BSD) or neomycin/kanamycin phosphotransferase (NeoR). mTagRFP was co-expressed with BSD to indicate chromosomal integration of donor DNA. On day 3 post-nucleofection, Blasticidin or G418 was added into the medium to select cells with integration. Resistant cells were amplified, and then transfected with Cre expression plasmid DNA to excise the fragment between loxP sites. After excision, Nodal open reading frame (ORF) is reconstituted with a loxP site and in-frame mCitrine. The residual loxP sequence is upstream of the Furin consensus motif and will be removed from the mCitrine::Nodal (mature) fusion by the convertase. Single cell FACS was performed to isolate successfully modified mono-clones. (B) Confirmation of integration. Three pairs of primers were used to genotype retrieved clones. Reaction (RXN) 1 with a pair of primers covers the entire integrated fragment, distinguishing modified allele from wide type allele via size difference. RNX2 and RNX3 amplify 5' and 3' adjacent regions, respectively. The entire integrated fragment was sequenced, and the adjacent regions are shown.

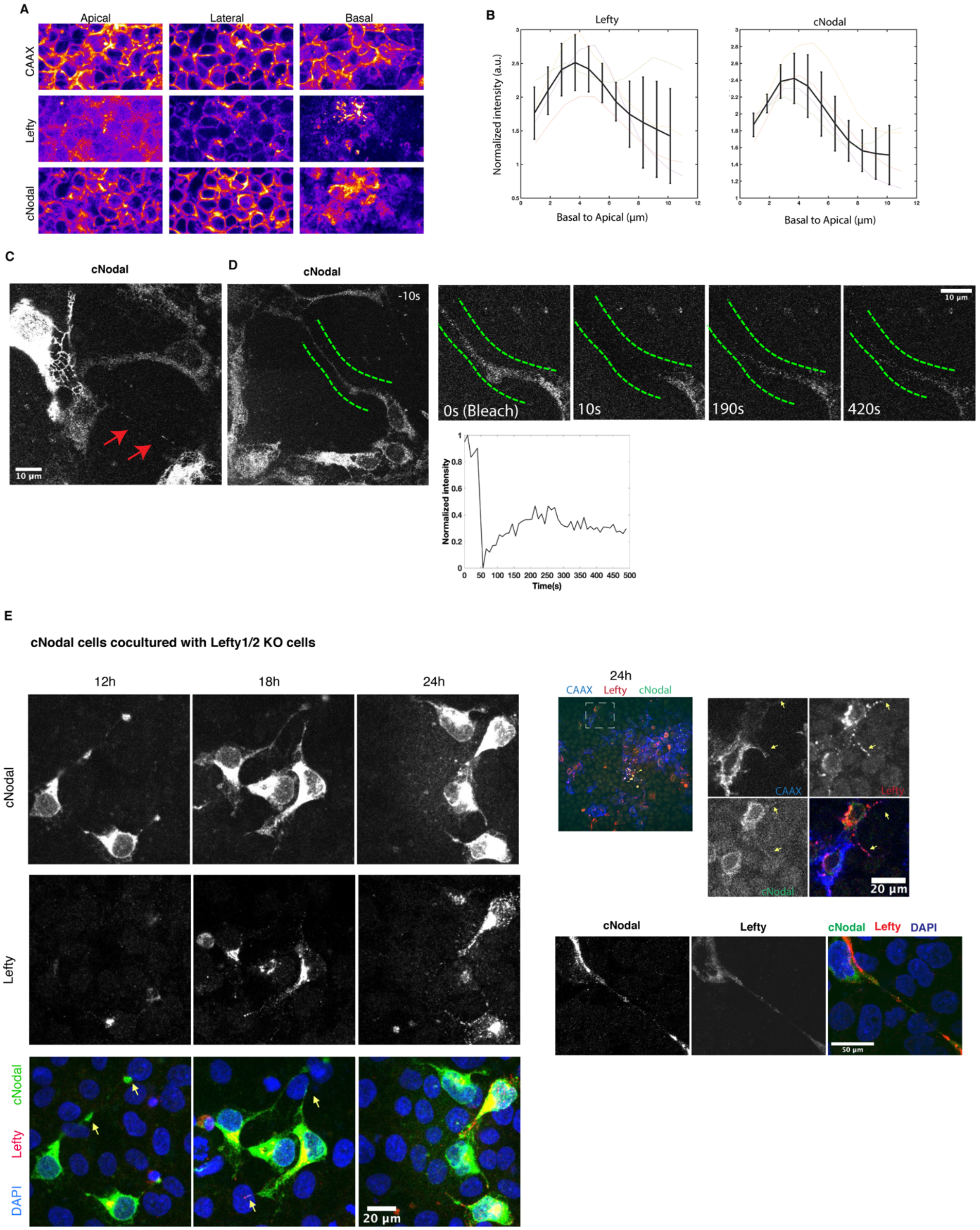

A
Apical
Lateral
Basal
CAAX
Lefty
cNodal
B
Lefty
cNodal
Normalized intensity (a.u.)
Basal to Apical (μm)
Basal to Apical (μm)
C
cNodal
10 μm
D
cNodal
-10s
0s (Bleach)
10s
190s
420s
10 μm
Normalized intensity
Time(s)
E
cNodal cells cocultured with Lefty1/2 KO cells
12h
18h
24h
cNodal
Lefty
cNodal
Lefty
DAPI
20 μm
24h
CAAX Lefty cNodal
CAAX
Lefty
cNodal
20 μm
cNodal
Lefty
cNodal Lefty DAPI
50 μm

**Supplementary figure 2. Nodal and Lefty proteins are found on the basal-lateral side of cells and in membrane protrusions.** (A-B) Immunostaining of cNodal (anti-GFP antibody) and Lefty (anti-Lefty) in Activin treated $Nodal^{cit/cit}$ cells with mCherry-CAAX membrane marker. Both cNodal and Lefty showed the highest levels at basal side (normalized to CAAX). (C) High magnification of cNodal live-cell imaging shows cNodal localized on thread-like structures connecting two cells. Red arrowheads indicate the cNodal puncta. (D) Fluorescence recovery after photobleaching (FRAP) shows cNodal was transported through a membrane protrusion connecting two cells. Green dash lines flank and indicate the photobleached protrusion. (E) Nodal and Lefty locate on CAAX labeled protrusions. Homozygous cNodal cells (mCherry-CAAX) were co-cultured with Lefty 1/2 compound knock-out cells and treated with Activin for indicated time. cNodal (anti-GFP antibody) and Lefty (anti-Lefty) were examined via immunostaining.

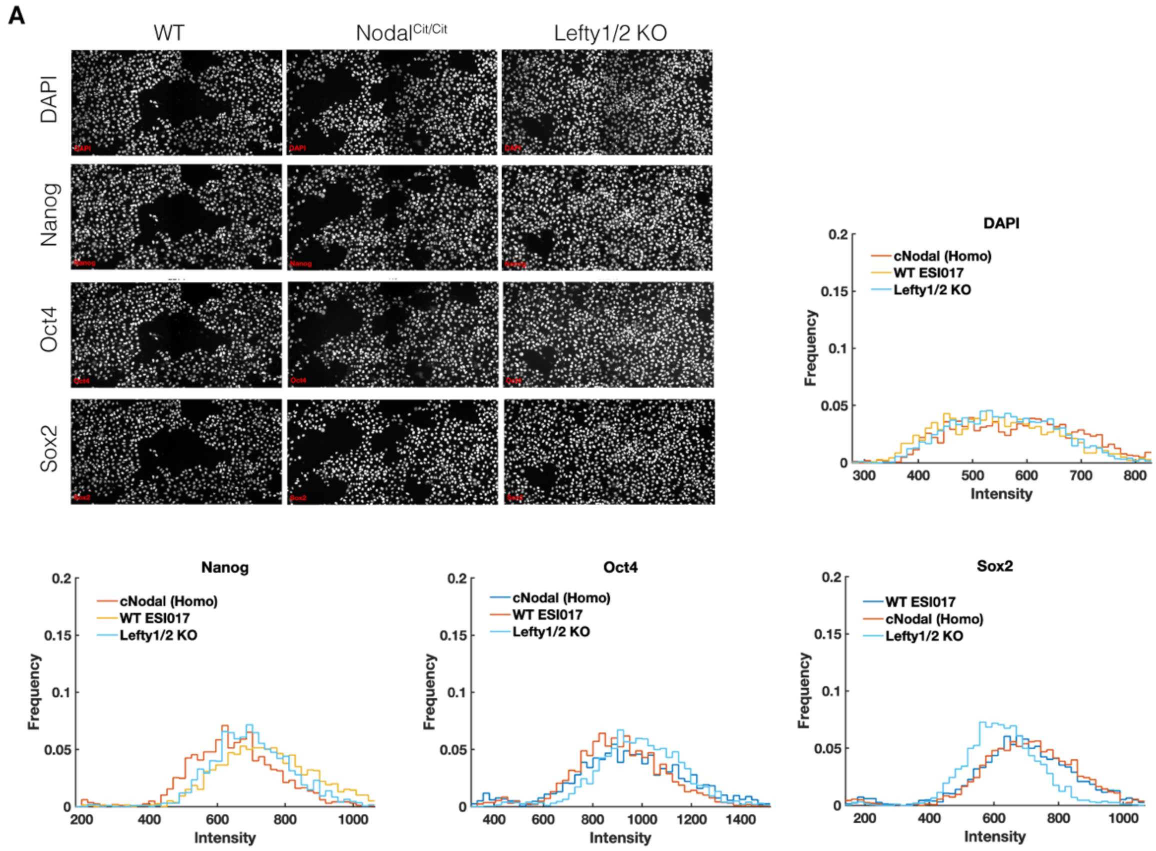


**Supplementary figure 3. cNodal knock-in and Lefty compound knock-out hESCs are pluripotent.** Pluripotency markers Oct4, Sox2 and Nanog levels were checked via immunofluorescence staining and compared to parental cells. WT, wild type ESI-017 cells. Nodal$^{\mathbf{cit/cit}}$ (clone H3), ESI-017 cells with biallelic mCitrine knock-in to Nodal locus. Lefty1/2 KO, ESI-017 cells with lefty 1 and Lefty 2 compound knock-out. DAPI was used to stain nuclei, Oct4, Sox2, Nanog levels are quantified based on fluorescence intensity.

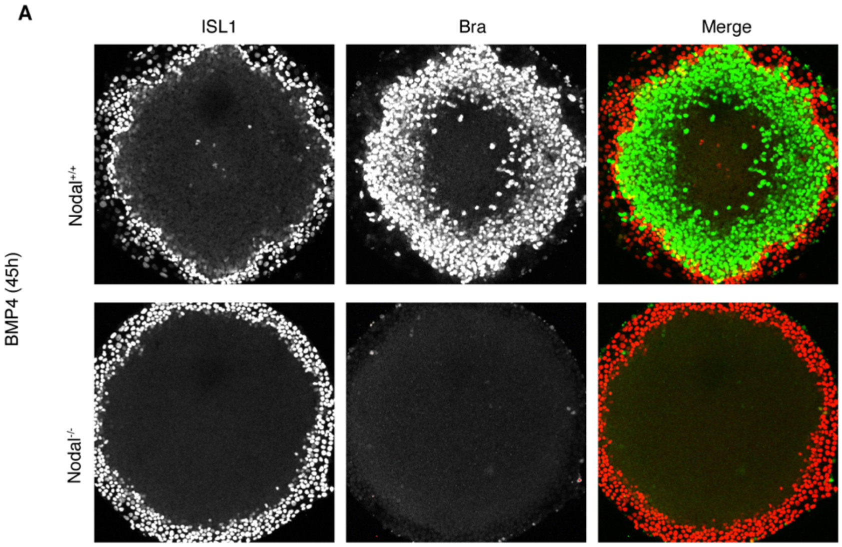


**Supplementary figure 4. Endogenous Nodal is required for mesodermal differentiation.** (A) Wide type ESI-017 ($Nodal^{+/+}$) or homozygous Nodal knockout ESI-017 ($Nodal^{-/-}$) cells were used to make micropatterned gastruloids, respectively. Extraembryonic fate maker ISL1 and Mesodermal fate maker Brachyury (BRA) are shown.

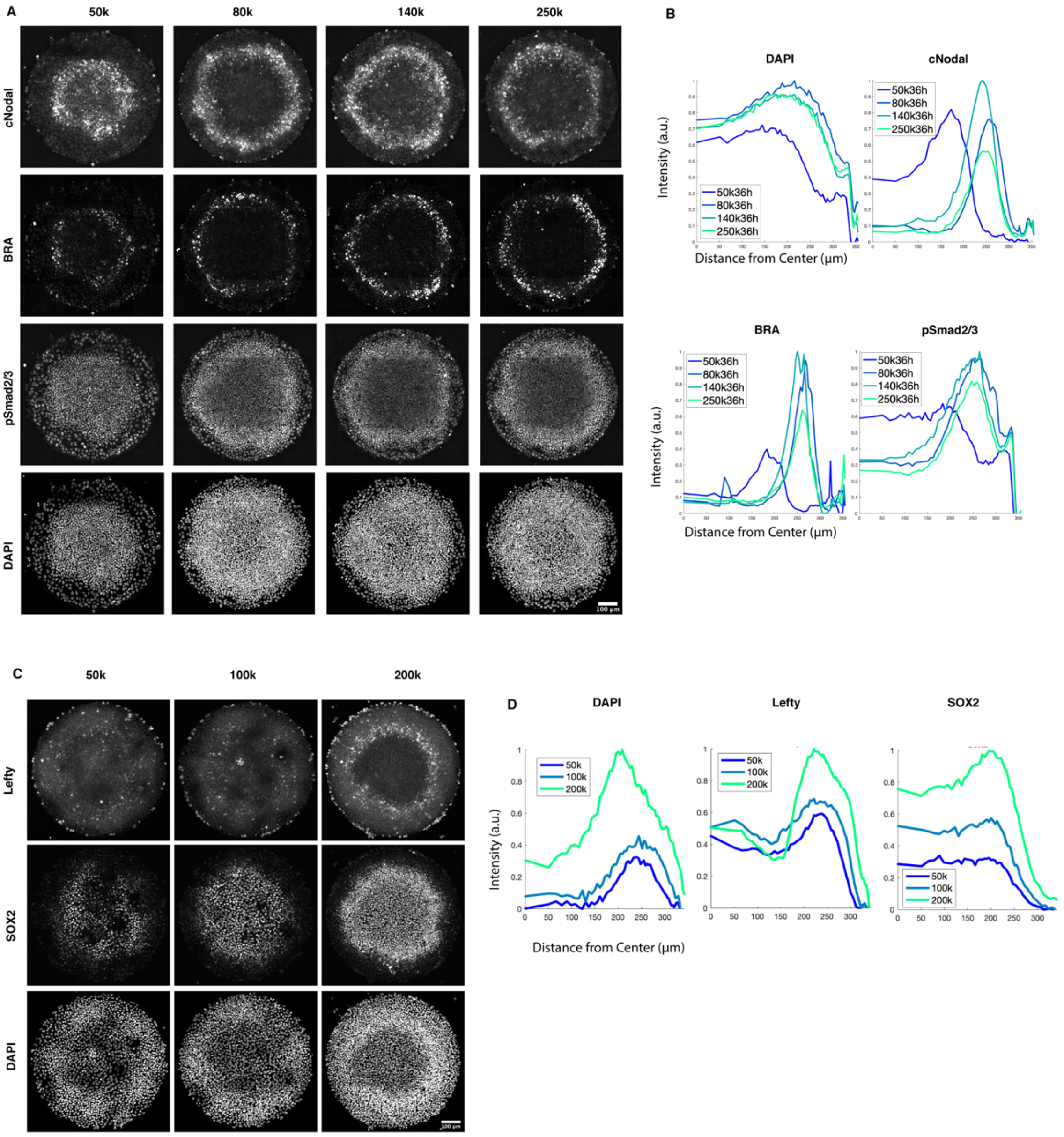


**Supplementary figure 5. Initial seeding cell density effects on Nodal and Lefty distribution in micropatterned gastruloids.** (A) Representative images of cNodal and pSmad2/3 distribution at different initial cell density conditions. 50,000, 80,000, 140,000, or 250,000 cells were initially seeded in each well of 96-well micropattern plate, respectively. Gastruloids were fixed at about39h post-BMP4 induction. (B) cNodal and pSmad2/3 levels were quantified as function of distance from colony center (n=6). (C) Representative images of Lefty protein and ectoderm/pluripotency marker Sox2 distribution at different initial cell density conditions. 50,000, 100,000, or 220,000 cells were initially seeded in each well of 96-well micropattern plate, respectively. Gastruloids were fixed at about39h post-BMP4 induction. (D) Lefty and Sox2 levels were quantified as function of distance from colony center (n=3).

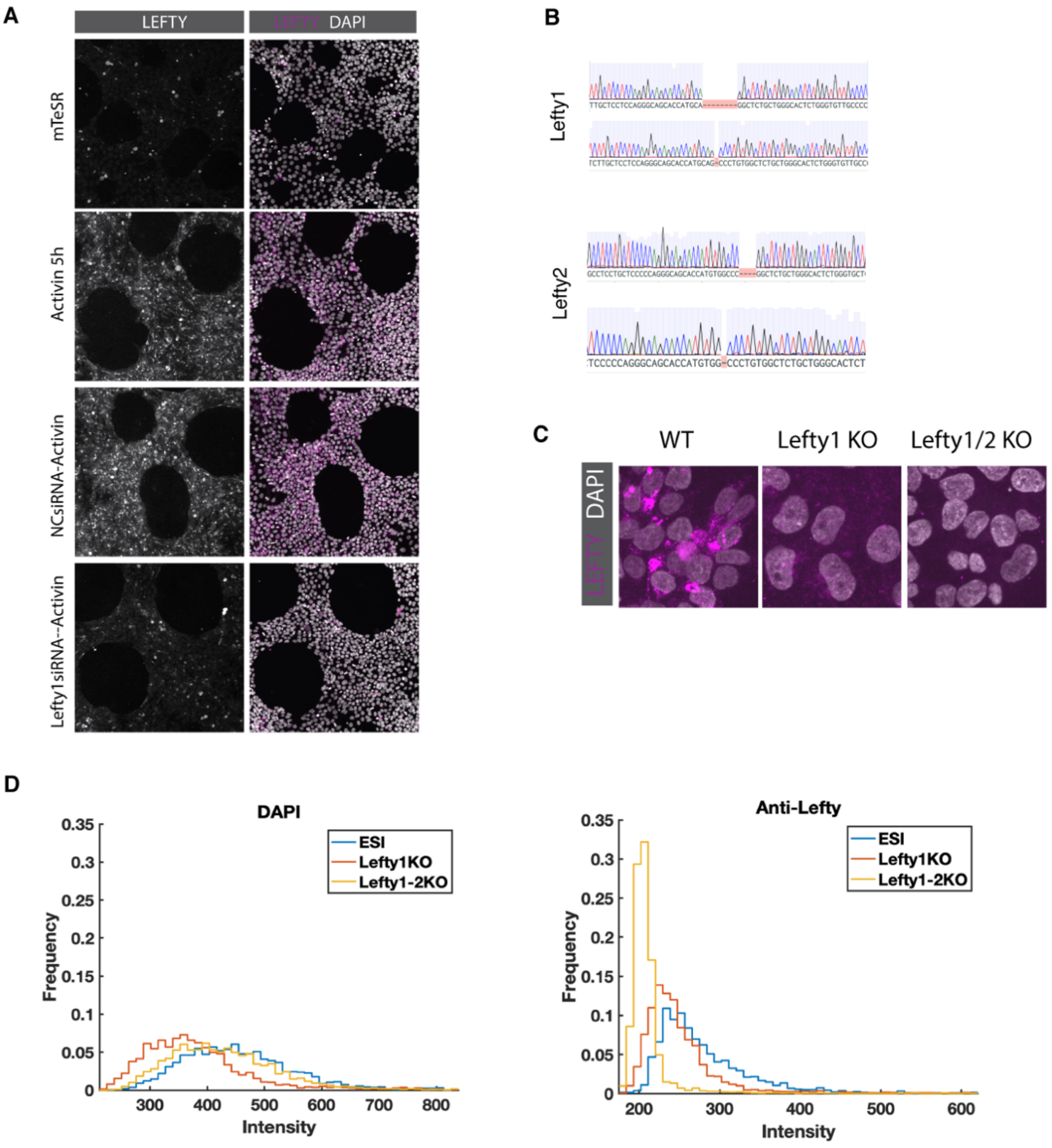


**Supplementary figure 6. Validation of Lefty1/2 knock-out.** (A) Lefty knock down experiments show that Lefty antibody staining quantitatively reflects the changing of Lefty expression levels. (B) Lefty 1/2 compound knock-out was verified by sequencing. Deletions causing open reading frame (ORF) shifts were found closely down-stream of the start codons of both alleles of the Lefty 1 and Lefty 2 genes. (C-D) Immunostaining using an antibody that recognizes both Lefty 1 and Lefty 2 confirmed ablation of Lefty proteins. (C) Representative images of ESI-017 cells, Lefty1 single knock-out or Lefty 1/2 compound knock-out cells treated with Activin and fixed. (D) Quantification of fluorescence intensity.

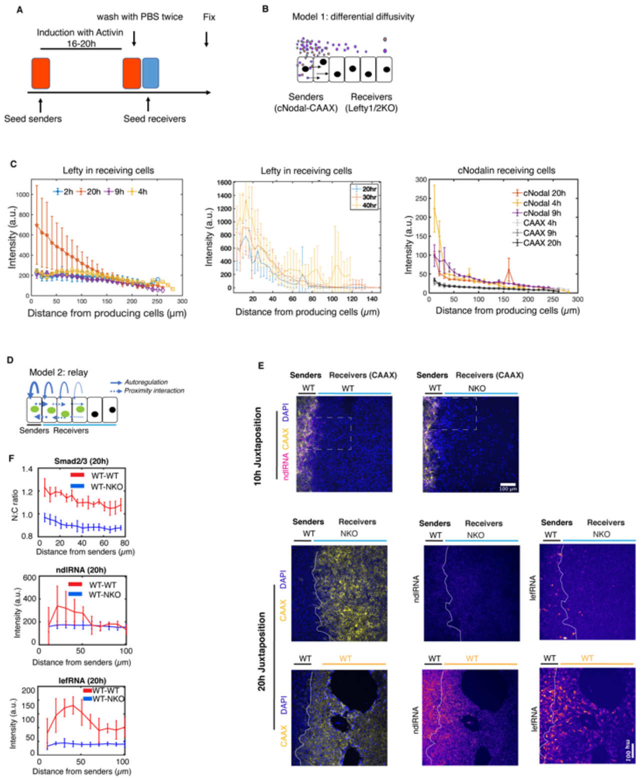

**Supplementary figure 7. Lefty protein is biochemically able to diffuse over several cell diamters while Nodal protein only reaches to immediately adjacent cells.** (A) Schematic illustration of juxtaposition experiments. In particular, sender cells (homozygous cNodal cells with mCherry-CAAX membrane label), were seeded in a well of culture insert, and then treated with 10 ng/ml Activin to induce expression of cNodal and Lefty. After 16-20h, induction medium was removed, and the sender cells were washed with PBS. The receiver Lefty1/2 compound knock-out or Nodal knock-out cells were then seeded juxtaposed to induced sender cells. (B) While sender cells produce and secrete cNodal and Lefty proteins, receiving cells cannot make either cNodal or Lefty. (C) Cells were fixed at various time points post-juxtaposition. Lefty and cNodal levels among receiving cells were examined through immunofluorescence staining and fluorescence intensity was quantified as function of distance from border of producing cells at indicated time. mCherry-CAAX that labels cell membrane of sender cells was also quantified as a control. (D) Schematic of relay model. Senders represent induced cells that express Nodal. Solid line with arrowhead, represents Nodal positive autoregulation. Dashed lines with arrowheads represent juxtacrine induction by Nodal. Back solid cycle represent nucleus, green solid cycle represents nuclear accumulated Smad2/3. (E-F) Experimental test of the relay model. (E) Nodal and Lefty RNA FISH to measure Nodal and Lefty transcripts in juxtaposition experiments with cells with the indicated genotypes. Receiver cells in all conditions were labelled by mCherry-CAAX. Upper panel shows an experiment in which the cells were fixed at 10h post-juxtaposition. The dashed boxes indicate areas shown in Figure 2F. (F) Experiments performed as in (E) and Nodal mRNA, Lefty mRNA and Smad2/3 were measured. Smad2/3 nuclear to cytoplasmic ratio, Nodal RNA and Lefty RNA are quantified as a function of distance from border (n=4 for each condition). Smad2/3 was measured in a separate experiment from Nodal and Lefty mRNA.

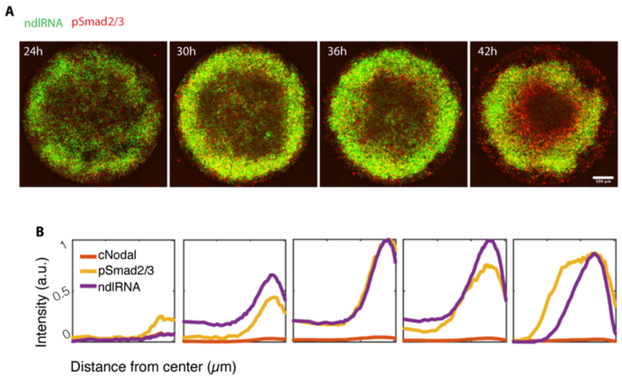


**Supplementary figure 8. Measurements of Nodal mRNA and pSmad2/3 in gastruloids with wildtype cells.** (A) Representative images of Nodal smFISH (green) and pSmad2/3 (red) at 42h of Nodal$^{+/+}$ without mCitrine tagging. Scale bar 100 µm. (B) Spontaneously quantification of Nodal RNA and protein, and pSmad2/3. Mean intensity is quantified over time as function of distance from colony center (n=6).

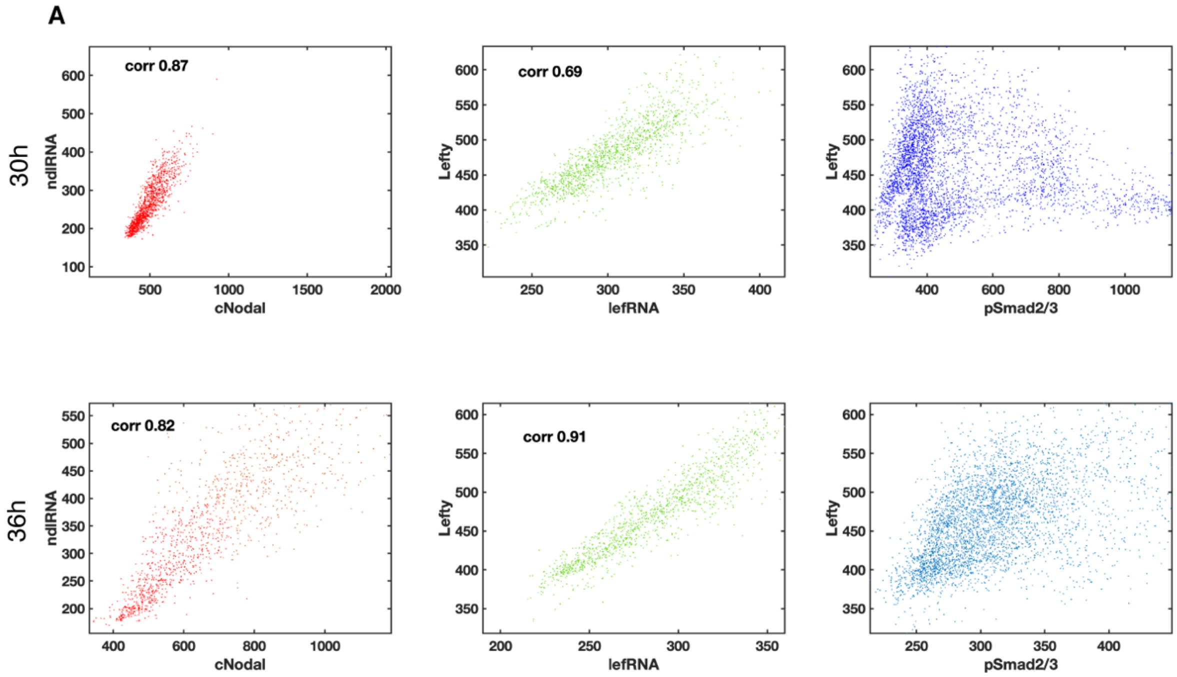


**Supplementary figure 9. Highly correlated mRNA and protein of cNodal and Lefty in gastruloids at 30h and 36h suggesting limited diffusion of the proteins from the sources.** mRNA and protein of cNodal and Lefty and pSmad2/3 in gastruloids at 30h and 36h were examined. The correlation of cNodal protein and mRNA (left-most panels), Lefty protein and mRNA (middle panels) levels in single cells were analyzed, correlation coefficient is shown. The relationship between Lefty protein and pSmad2/3 was also analyzed.

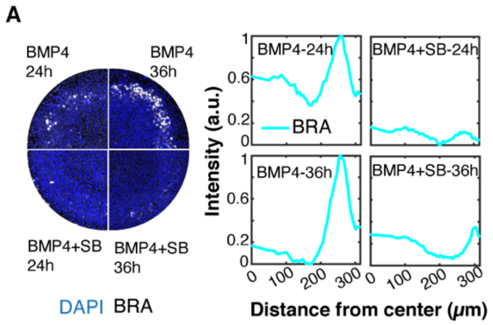


**Supplementary figure 10. Position of mesodermal domain is set at early stage and maintains.** (A) Representative images of mesodermal fate maker Brachyury (BRA) at 24h or 36h post-BMP4 treatment, with or without addition of Nodal signaling inhibitor SB-431542. (B) The levels of BRA were quantified as function of distance from colony center. Normalization was performed so that at each time point the maximum value of both curves is 1 and the minimum value is 0 (n=6).

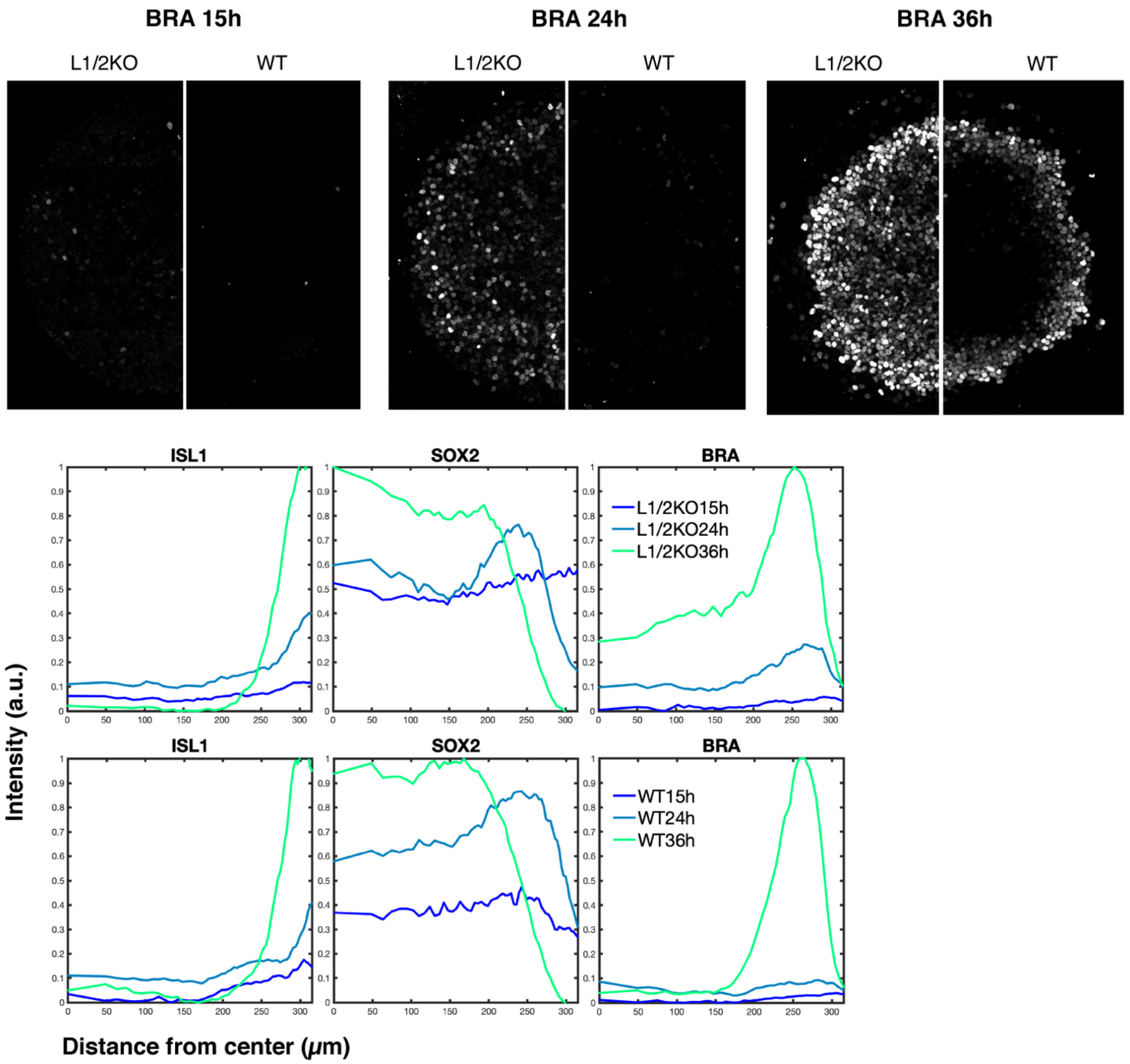


**Supplementary figure 11. Lefty1/2 knock-out causes earlier and broader expression of mesodermal marker Brachyury.** Gastruloids created with wild type or Lefty 1/2 compound knock-out cells were fixed at 15h, 24h, and 36h, respectively. Extraembryonic marker ISL1, mesodermal marker Brachyury (BRA), and ectodermal/pluripotency marker Sox2 were examined by immunofluorescence staining. Representative images of BRA at indicated time points are shown in upper panel, mean intensity of ISL1, Sox2 and BRA as function of distance from colony center is shown in lower panel (n=6).

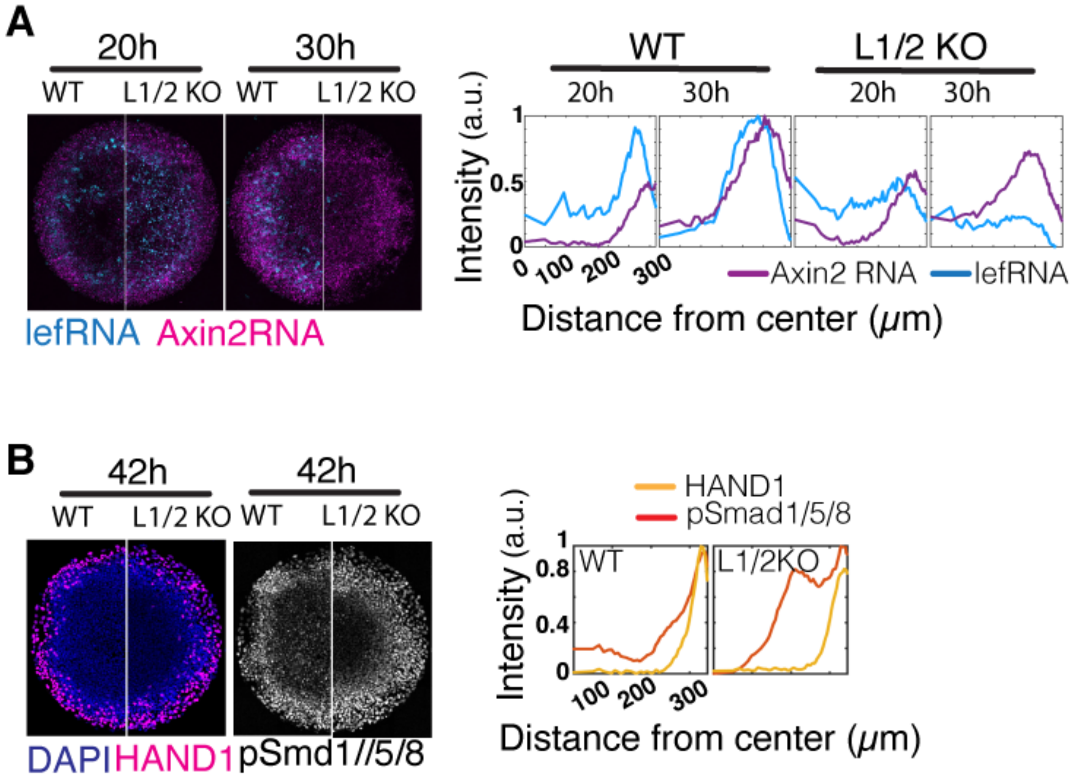


**Supplementary figure 12. Lefty1/2 knock-out leads to subtle changes in BMP4 and Wnt signaling.** (A) Analysis of Wnt signaling. Transcripts of Axin2, a Wnt signaling transcriptional target, and transcripts of Lefty1/2 in gastruloids at 20h and 30h were co-examined via smFISH. Mean intensity of FISH signal was quantified as function of distance from colony center (n=6). (B) Analysis of BMP signaling. HAND1, a BMP4 target gene, pSmad1/5/8, BMP signaling transducers, in gastruloids at 42h were examined, respectively. Mean intensity is quantified as a function of distance from colony center [illegible]

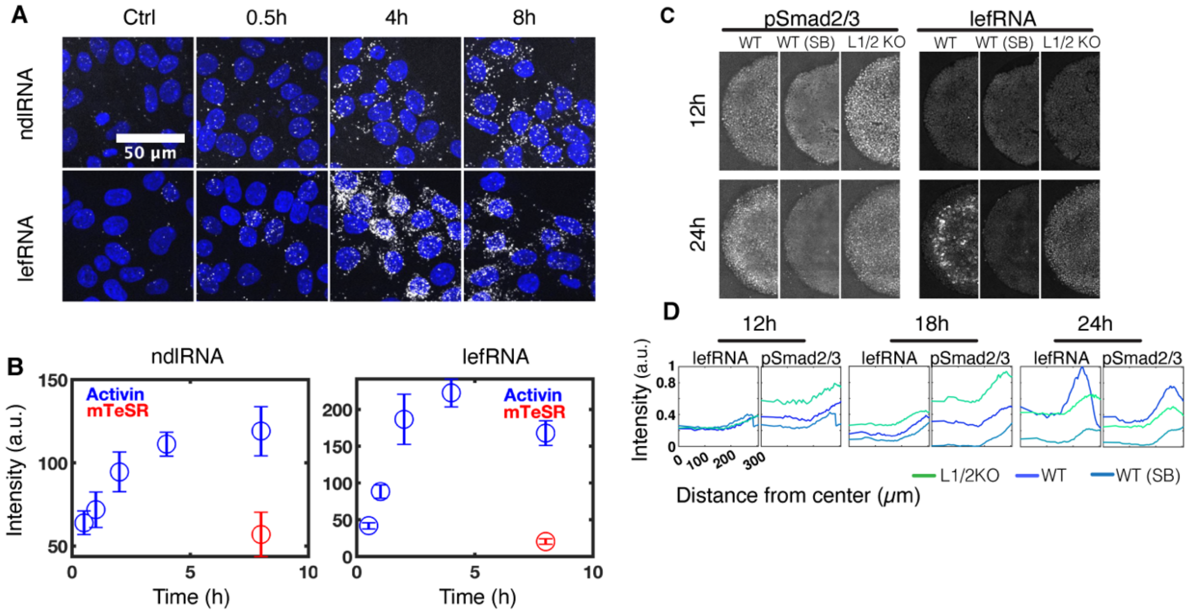


**Supplementary figure 13. smFISH shows dynamics of Lefty transcription in standard culture and gastruloids** (A-B) smFISH of Nodal and Lefty RNA. Cells were treated with Activin for 0.5h, 4h, 8h or no treatment for 8h. (A) Representative images of Nodal smFISH (upper panel), Lefty smFISH (lower panel), nuclei were indicated by DAPI (blue), scale bar 50 µm. (B) quantification of smFISH signal in each nucleus, mean value of four images for each condition is shown, SEM is shown. (C-D) Lefty transcription (lefRNA) and pSmad2/3 in WT or Lefty1/2 KO gastruloids. (C) Representative images of Lefty smFISH and pSmad2/3 staining at 12h and 24h of gastruloids made with WT hESCs, Lefty1/2 compound knockout (L1/2 KO) cells or WT hESCs in the presence of SB-431542 are shown. (D) Mean intensity of signal of interest at 12h, 18h and 24h was quantified as function of distance from center, n=6.
